# Accelerated DNA evolution in rats is driven by differential methylation in sperm

**DOI:** 10.1101/033571

**Authors:** Xiao-Hui Liu, Jin-Min Lian, Fei Ling, Ning Li, Da-Wei Wang, Ying Song, Qi-Ye Li, Ya-Bin Jin, Zhi-Yong Feng, Lin Cong, Dan-Dan Yao, Jing-Jing Sui

## Abstract

Lamarckian inheritance has been largely discredited until the recent discovery of transgenerational epigenetic inheritance. However, transgenerational epigenetic inheritance is still under debate for unable to rule out DNA sequence changes as the underlying cause for heritability. Here, through profiling of the sperm methylomes and genomes of two recently diverged rat subspecies, we analyzed the relationship between epigenetic variation and DNA variation, and their relative contribution to evolution of species. We found that only epigenetic markers located in differentially methylated regions (DMRs) between subspecies, but not within subspecies, can be stably and effectively passed through generations. DMRs in response to both random and stable environmental difference show increased nucleotide diversity, and we demonstrated that it is variance of methylation level but not deamination caused by methylation driving increasing of nucleotide diversity in DMRs, indicating strong relationship between environment-associated changes of chromatin accessibility and increased nucleotide diversity. Further, we detected that accelerated fixation of DNA variants occur only in inter-subspecies DMRs in response to stable environmental difference but not intra-subspecies DMRs in response to random environmental difference or non-DMRs, indicating that this process is possibly driven by environment-associated fixation of divergent methylation status. Our results thus establish a bridge between Lamarckian inheritance and Darwinian selection.

---

DNA variation passed stably from parent to offspring is the traditional mechanism underlying trait heritability, and provides the basis for Darwinian selection. In contrast, transgenerational inheritance of epigenetic variation has recently been proposed as a form of Lamarckian acquired inheritance, where species adapt to the changing environment without an accompanying DNA sequence change (Van Soom et al. 2014). Transgenerational epigenetic inheritance has been controversial as it was previously thought that the epigenome is fully erased and reestablished between generations in order for appropriate cellular development and differentiation to occur in mammals (Daxinger and Whitelaw 2012; Franklin and Mansuy 2010; Heard and Martienssen 2014). In recent years, Lamarckian acquired inheritance has gained increasing support due to the findings of transgenerational transmission of epigenetic status in a number of species (Daxinger and Whitelaw 2012; Franklin and Mansuy 2010; Heard and Martienssen 2014; Lim and Brunet 2013; Van Soom et al. 2014). However, transgenerational epigenetic inheritance and Lamarckian acquired inheritance is still under debate for unable to rule out DNA sequence changes as the underlying cause for heritability (Heard and Martienssen 2014; Lim and Brunet 2013). Distinguishing the relative contributions of epigenetic changes and DNA variation to phenotypic variation and determining the potential for epigenetics to impact DNA evolution are keys to resolving these questions (Boffelli and Martin 2012; Heard and Martienssen 2014; Lim and Brunet 2013). The mutation rate at CpG sites is influenced by methylation status (Fryxell and Moon 2005; Mugal and Ellegren 2011; Xia et al. 2012; Zhao and Jiang 2007), making methylation a plausible mechanism by which the epigenome influences heritable traits through changes to the underlying DNA sequence. However, the relationship between epigenetic variation and DNA variation, their relative contribution to phenotypic divergence, and relative impact on the evolution of species are far from clear (Heard and Martienssen 2014; Lim and Brunet 2013).

The Norway rat originated in South China 1.2-1.6 million years ago and spread throughout the rest of the world with humans (Song et al. 2014; Wu and Wang 2012). The split of two subspecies *Rattus norvegicus caraco* (Rnc) and *Rattus norvegicus norvegicus (Run)*, distributed in North and South China respectively, has been supported by both morphological data and mitochondrial DNA analysis (Song et al. 2014). *R. n. norvegicus* breed year round while *R. n. caraco* has restricted breeding during the winter (Wang et al. 2011). Intriguingly, we observed that the latter can breed year round when reared in proper room conditions, indicating the reproductive activity of the Norway rat is very sensitive to environmental change. The plasticity of this trait may indicate that it is an example of Lamarckian inheritance. Additionally, reproductive behavior involves the coordinated expression of a cohort of genes, making it likely that epigenetic inheritance could play a role. Thus, these two subspecies provide a nice model to explore the relationship between epigenetic variation and DNA variation and their contribution to species divergence. In this study, we analyzed DNA sequence and methylation variation in sperm to illustrate the relationship between transgenerational epigenetic inheritance and DNA evolution in the divergence of *R. n. norvegicus* and *R. n. caraco*. We found that all kinds of environment-associated methylation differences can lead to increasing of nucleotide diversity, and fixation of methylation status can further lead to accelerated fixation of DNA variants. These results establish a bridge between Lamarckian acquired inheritance and Darwinian selection.

## Results

### DNA evolution is associated with environment-dependent methylation pattern

Two individuals were selected from each Norway rat subspecies *R. n. caraco* and *R. n. norvegicus*, located in Harbin City (126°32′E, 45°48′N) and Zhanjiang City (110°21′E, 21°16′N), respectively. Both the genome and sperm methylome were sequenced for each sample, and each methylome was proofread using the corresponding genome sequence of this individual. On average, we generated more than 20× coverage of genome sequence for each sample, and over 89% of the non-gap genome was covered by more than one read (Supplemental Table 1). The basic BS-seq data analysis was conducted using a custom pipeline validated in several previous projects (Bonasio et al. 2012; Shao et al. 2014). Each methylome was proofread by genome sequence of this individual. About 22 million CpGs (90% of all rat CpGs) were covered by at least one read. The average read coverage for CpGs is at least 23× per sample with an overall methylation level of 76% for all CpG sites (Supplemental Table 1). The non-CpG sites were not significantly methylated in any of the datasets.

In total, we obtained 1,859 inter-subspecies DMRs (Differentially methylated regions) with a total sequence length of 2.60 Mb (Supplemental Table 2), 2,669 intra-*Rnn* DMRs with a total length of 2.67 Mb (Supplemental Table 3), and 3,461 intra-*Rnc* DMRs with a length of 3.46 Mb (Supplemental Table 4). We found that in 87.74% of inter-subspecies DMRs methylation levels are the same between individuals within each subspecies (Supplemental Fig. 1A), indicating the methylation statuses in the inter-subspecies DMRs are subspecies specific and retained across generations. Although the total sequence length of each DMR set covers only one thousandth of the genome, each set of DMRs was distributed evenly throughout the genome (Supplemental Fig. 1B). Genome sequences were classified as DMRs or non-DMRs for further analysis.

As Figure 1 illustrates, phylogenetic analysis of sperm methylomes, genome-wide single nucleotide variants (SNVs), DMR sequences and sampled non-DMR sequence (the first sampling method) all support the divergence between the two subspecies, but show different tree topologies. The topology of the tree built using DNA sequences in inter-subspecies DMRs is similar to the topology of the clustering tree constructed using methylation patterns (Fig. 1A, 1B and 1D) but not sampled non-DMR sequences or genome-wide SNVs, and the topology of the tree built using sampled non-DMR sequences is similar to the topology of the tree constructed using genome-wide SNVs (Fig. 1C and 1E), indicating a possible correlation between methylation pattern and DNA divergence. Compared to sampled non-DMR sequence with equal length and GC content, DNA sequences in inter-subspecies DMRs show increased mean distance between subspecies (independent t test: *p* = 0; Fig. 1; Supplemental Fig. 2A), but decreased mean distance within subspecies (independent t test: *p* < 0.01; Fig. 1; Supplemental Fig. 2B and 2C), indicating that methylation variation in inter-subspecies DMRs may promote species divergence while decreases the within-species DNA divergence.

**Figure 1.**
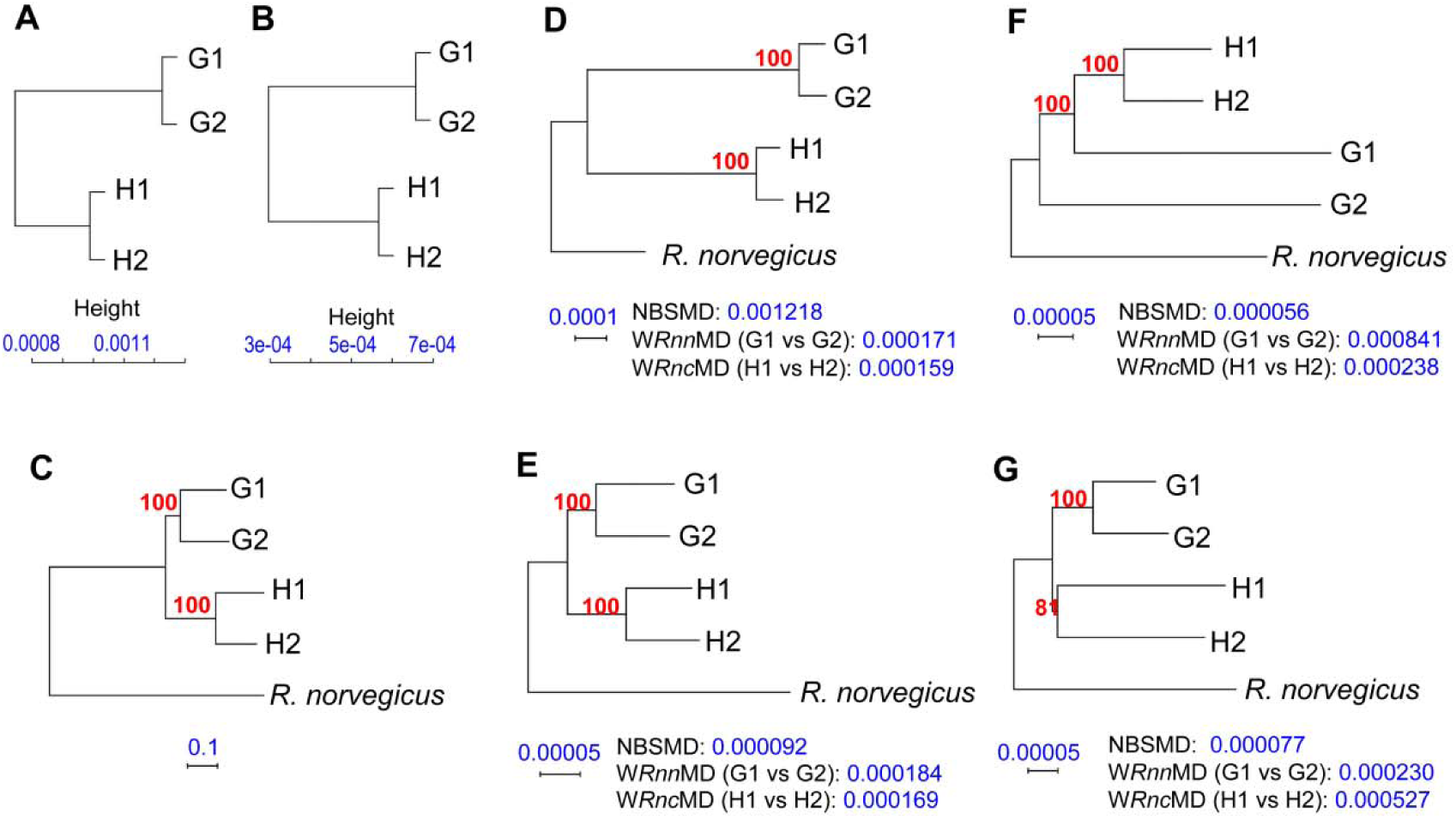
Evolution patterns of inter-subspecies DMRs, intra-subspecies DMRs and non-DMRs. Rat G1 and G2 are two individuals of *R. n. norvegicus*, and Rat H1 and H2 are two individuals of *R. n. caraco*. Non-DMRs were sampled using the first method with equal length and similar GC content as inter-subspecies DMRs. Except methylation trees, all trees were constructed using the Neighbor-Joining method in the PHYLIP software package. **NBSMD**: Net between subspecies mean distance. **W*Rnn*MD**: Within *Rnn* mean distance. **W*Rnc*MD**: Within *Rnc* mean distance. (*A*) Methylation clustering tree for the 4 rats. Methylation levels are calculated using genes. *(B)* Methylation clustering tree for the 4 rats. Methylation levels are calculated using 10kb sliding windows. (*C*) Phylogenetic tree constructed using genome-wide SNVs. (D) The phylogenetic tree constructed using combined inter-subspecies DMR sequences. *(E)* The phylogenetic tree constructed using a set of sampled non-DMR sequences. *(F)* The phylogenetic tree constructed using combined intra-*Rnn* DMR sequences. *(G)* The phylogenetic tree constructed using combined intra-*Rnc* DMR sequences.

In comparison, both phylogenetic trees built with the DNA sequences in intra-subspecies DMRs have decreased mean distances between subspecies than both inter-subspecies DMRs and non-DMRs (Fig. 1F and 1G). The phylogenetic tree built using four sequences in intra-*Rnn* DMRs shows increased distance within *Rnn* but not *Rnc* (Fig. 1F), and the phylogenetic tree built with the four sequences in intra-*Rnc* DMRs shows increased distance within *Rnc* but not *Rnn* (Fig. 1G). This indicates that methylation variation and its related DNA variation in intra-subspecies DMRs are not associated with species divergence.

We calculated Tajima’s D to test for departure from neutrality in these regions. Tajima’s D is 1.678528, -0.393469 and -0.277028 in combined inter-subspecies, intra-*Rnn* and intra-*Rnc* DMR sequences, respectively (Fig. 2). In comparison, the mean Tajima’s D in 1,000 sampled non-DMR sequences is 0.002920 ± 0.099323 (SD), with a maximum value of 0.453000, which is significantly lower than in inter-subspecies DMRs (independent t test: *p* = 0; Fig. 2). The distribution of Tajima’s D in non-DMRs is normal with a median value of 0, indicating the sampled non-DMRs are a set of random neutral sequences. The increased Tajima’s D value in the inter-subspecies DMRs indicates strong selection and existence of a larger proportion of fixed or fixing SNVs between subspecies in inter-subspecies DMRs. The significantly negative Tajima’s D in both sets of intra-subspecies DMRs indicates an excess of random substitutions (Fig. 2).

**Figure 2.**
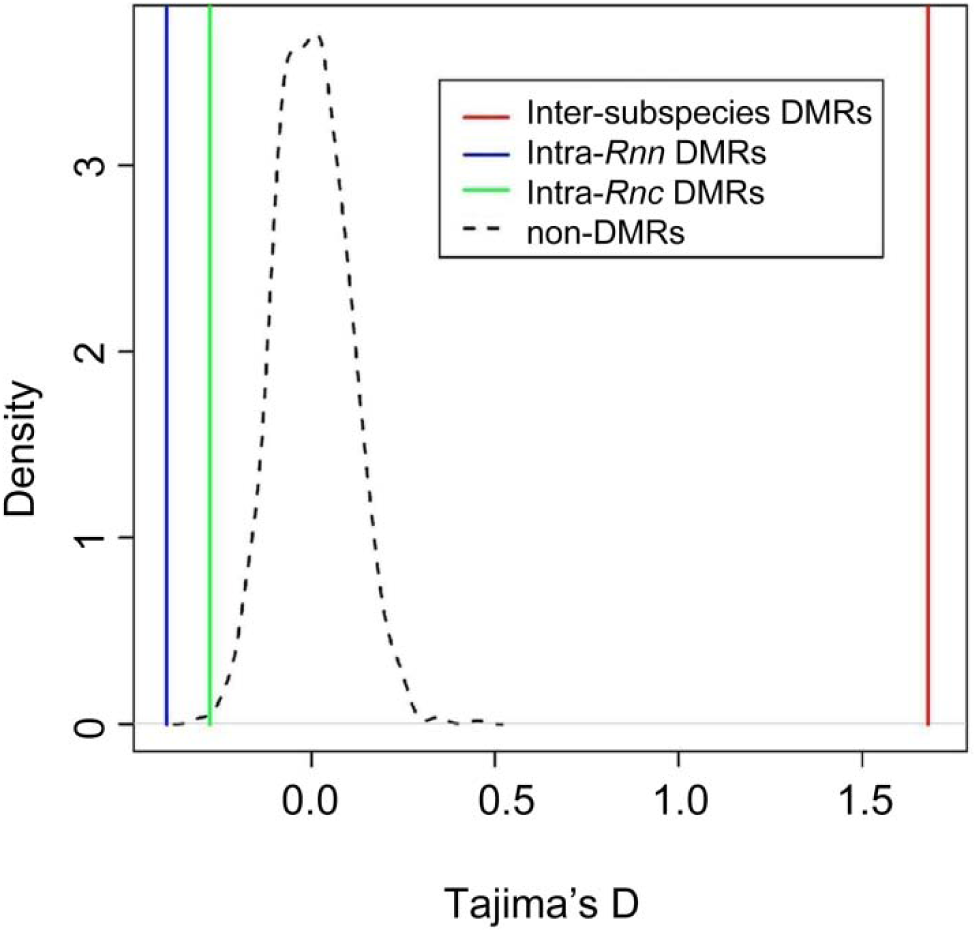
Comparison of Tajima’s D using the second sampling method.

We wanted to characterize SNVs that are fixed or fixing in one of the subspecies, which we will refer to as subspecies specific SNVs (SS-SNVs) (Supplemental Fig. 3), To accomplish this we compared the DNA divergence level between DMRs and non-DMRs by calculating the ratio of SS-SNVs (RSS, SS-SNVs were identified under *p* value of 0.05 and 0.01, respectively) using 10 additional genome sequences (5 for each subspecies, kindly provided by the Kunming Institute of Zoology, CAS). Inter-subspecies DMRs have significantly higher RSSs than non-DMRs (Chi-Square test; DMRs: 42.95% at significant level < 0.05 of SS-SNVs statistic, 30.88% < 0.01; non-DMRs: 14.87% < 0.05, 8.87% < 0.01; Table 1; x^2^ = 1587.585, df = 1, *p* < 2.2e-16, under the p value of 0.05; x^2^ = 1786.16, df = 1, *p* < 2.2e-16, under the *p* value of 0.01). These results further confirmed the accelerated DNA evolution in inter-subspecies DMRs.

**Table 1.** Population test of RSS

| Location | SNVs<br>identified in 4<br>genomes | SS-SNVs identified<br>in 4 genomes (RSS) | SNVs covered<br>in 14 genomes | SS-SNVs identified<br>in 14 genomes ($p$<br>< 0.01) (RSS) | SS-SNVs identified<br>in 14 genomes ($p$ <<br>0.05) (RSS) |
| --- | --- | --- | --- | --- | --- |
| Genome | 12,669,132 | 2,722,049 | 3,236,117 | 28,7893 | 482,520 |
| non-DMRs | 12,648,498 | 2,711,624 (21.44%) | 3,231,817 | 286,565(8.87%) | 480,673(14.87%) |
| DMRs | 20,634 | 10,425 (50.52%) | 4,300 | 1,328(30.88%) | 1,847(42.95%) |

### Nucleotide diversity is promoted by variation of methylation level rather than methylation level itself

To explore the relationship between DNA methylation variation and substitution rate, we compared nucleotide diversity (π) between DMRs and non-DMRs. Using the second sample method of non-DMRs, we found that the distribution pattern of π was positively skewed in both inter- and intra-subspecies DMRs by some higher π values, but was normal in both inter- and intra-subspecies non-DMRs (Supplemental Fig. 4). Mann-Whitney tests revealed π in both inter- and intra-subspecies DMRs was significantly higher than that of non-DMRs *(p* < 0.00001; Fig. 3A, Supplemental Table 5). We also calculated π in the combined DMRs of the whole genome for the inter-subspecies, intra-*Rnn*, intra-*Rnc* DMRs and sampled non-DMRs (by the second sampling method), respectively. The distribution of π in the 1,000 sampled combined non-DMRs is normal with a small standard deviation (0.001844 ± 0.000053; Fig. 3B), indicating DNA variation patterns in non-DMRs are similar and random. The value of π in the combined inter-subspecies, intra-*Rnn* and intra-*Rnc* DMR sets is 0.00343, 0.003055 and 0.0026, respectively, which are significantly higher than both the average and the maximum value of sampled non-DMRs (independent t test: *p* = 0; Fig. 3B), indicating extremely high levels of DNA variation within DMRs. We then stratified DMRs by methylation level and found that DMRs have a significantly higher π than non-DMRs irrespective of the methylation level (Mann-Whitney test: *p* < 0.00001; Fig. 4, Supplemental Table 6), suggesting that the nucleotide diversity in DMRs could be determined by variation of methylation level rather than methylation level itself.

**Figure 3.**
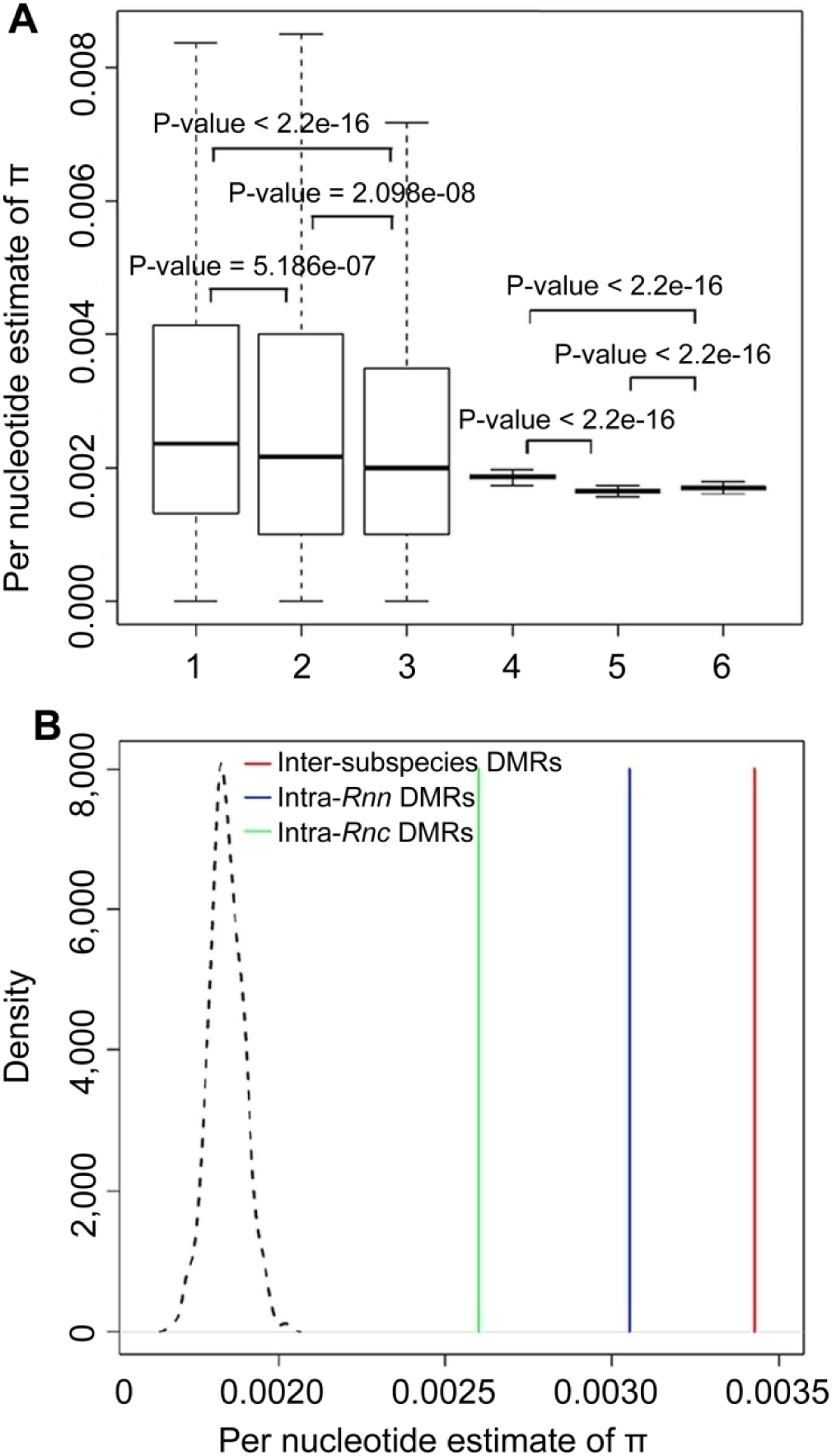
DMRs in response to both random and stable environmental difference show increased nucleotide diversity. (*A*) Comparison of π using second sampling method, π of inter-subspecies DMRs or non-DMRs are calculated using 4 individuals, and π of intra-subspecies DMRs or non-DMRs are calculated using 2 individuals of each subspecies respectively. Coordinates of horizontal axis: 1. Inter-subspecies DMRs; 2. Intra-*Rnn* DMRs: 3. Intra-*Rnc* DMRs: 4. Inter-subspecies non-DMRs; 5. Intra-*Rnn* non-DMRs; 6. Intra-*Rnc*-DMRs. Comparisons of π in each set are made using a Mann-Whitney test. (*B)* Comparison of π using the first sampling method.

**Figure 4.**
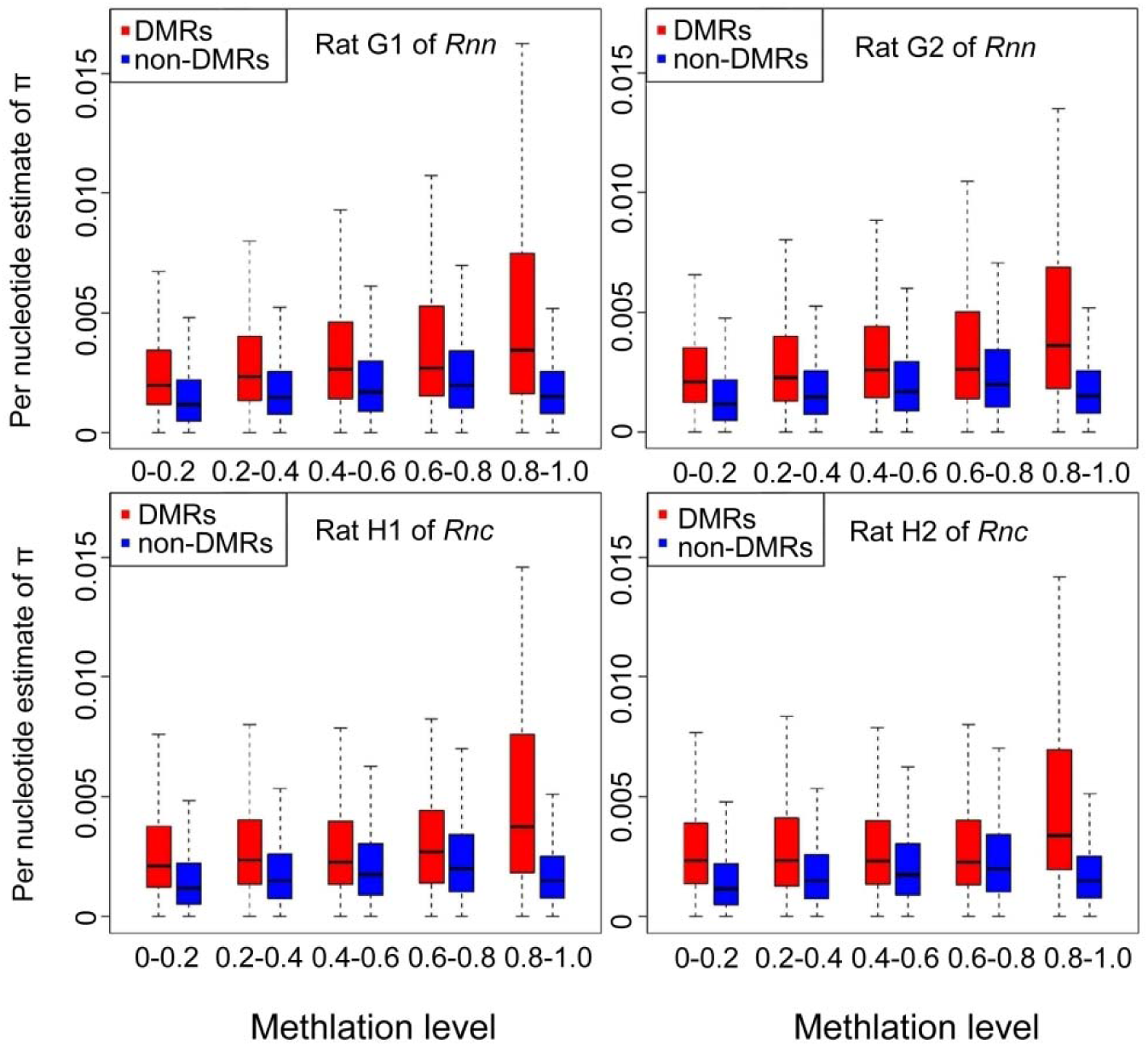
DMRs have a significantly higher π than non-DMRs irrespective of the methylation level. Here, π is calculated using all 4 individuals.

Deamination caused by methylation leads to increased rates of C to T substitution (Ehrlich et al. 1986; Jiang et al. 2007; Zhao and Jiang 2007). To examine the possible role of deamination on increased nucleotide diversity in DMRs, We first compared all six kinds of DNA substitutions (A/G, A/C, A/T, C/T, C/G and T/G) in inter-subspecies DMRs and non-DMRs, as well as across the whole genome after normalization. The substitution rates of C/T and A/G were significantly higher than the other four types of DNA substitutions (A/C, T/G, C/G and A/T) both in DMRs and non-DMRs (Independent T test: *p* < 0.0001; Fig. 5A) indicating the prevalence of deamination and the higher rate of transitions. However, the normalized rate of each substitution type in inter-subspecies DMRs is 2.5 times higher than that of non-DMRs and of the whole genome (Independent T test: *p* < 0.0001; Fig. 5A), suggesting that methylation variation affects all 6 nucleotide substitution types, not just types related to methylation.

**Figure 5.**
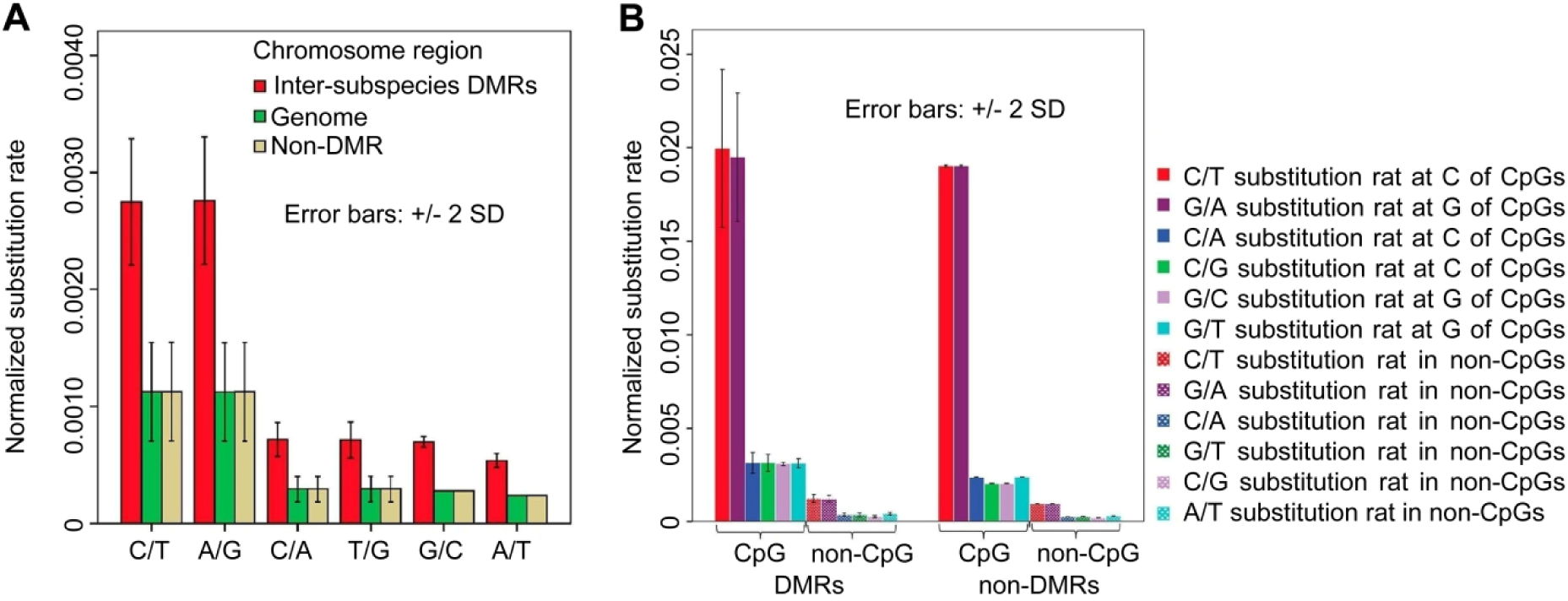
Deamination caused by methylation is not the root cause of increased nucleotide diversity in DMRs. (*A*) Comparison of 6 different types of DNA substitution rates between inter-subspecies DMRs and non-DMRs. *(B)* Comparison of different types of DNA substitution rate in CpG and non-CpG sites between inter-subspecies DMRs and non-DMRs using the second sampling method.

Because methylation and deamination mainly occur at the cytosine base in CpGs, the C/T substitution rate at these sites, as well as paired G/A on another strand, should be higher than that of non-CpGs. We compared normalized DNA substitution rates in CpG sites and non-CpG sites between DMRs and non-DMRs (using the second non-DMR sampling method to decrease the influence of sequence length on substitution rate calculation). As expected, the normalized rates of C/T and G/A substitutions in CpGs are higher than in non-CpGs in both DMRs and non-DMRs (Mann-Whitney test: *p* < 0.0001; Fig. 5B), which demonstrates the strong impact of deamination on C/T substitutions. We then compared DNA substitution rates in CpG sites and non-CpG sites between DMRs and non-DMRs, respectively. In non-CpG sites, all 6 normalized DNA substitutions are significantly higher in DMRs than in non-DMRs (Mann-Whitney test: *p* < 0.01; Fig. 5B). This result demonstrates that the difference in nucleotide diversity between DMRs and non-DMRs is not a consequence of deamination because non-CpG sites are basically unaffected by methylation. However, in CpG sites, neither C/T nor G/A substitution rates are significantly different between DMRs and non-DMRs (Mann-Whitney test: *p* > 0.05; Fig. 5B), whereas the other 4 types of substitution (C/A and C/G at cytosine of CpG; G/T and G/C at guanine of CpG) are significantly higher within DMRs compared to non-DMRs (Mann-Whitney test: *p* < 0.01; Fig. 5B). This result indicates that methylation-induced deamination cannot account for the increased rates of C/T and G/A substitutions in DMRs. Meanwhile, the increased rates of C/T and G/A substitutions in DMRs imply that deamination exerts relatively lower influence on C/T substitution at CpG sites in DMRs than in non-DMRs. In other words, CpG sites are relatively more conserved in DMRs than in non-DMRs. Thus, the above results support the idea that deamination caused by methylation is not the root cause of increased nucleotide diversity in DMRs, and that it is variance of methylation level but not methylation itself influencing the substitution rates of DMRs.

### Conserved CpG content in DMRs indicates additional forces drive methylation differences

DNA methylation is highly associated with the genomic and functional context (Gutierrez-Arcelus et al. 2013). We characterized CpG contents in DMRs to explore the possible influence of DNA variation on methylation patterns. The average CpG number per DMR is 27.03 ± 18.30 (average density = 0.0248 ± 0.0180) in inter-subspecies DMRs, 19.01 ± 7.73 (average density = 0.0190 ± 0.0073) in intra-*Rnn* DMRs, 19.95 ± 7.94 (average density = 0.0199 ± 0.0079) in intra-*Rnc* DMRs (Supplemental Fig. 5A). The average CpG density of combined DMRs (0.019430 ± 0.000478) is significantly higher than that of non-DMRs (0.012789 ± 0.000153) (Mann-Whitney test: *p* < 0.00001) (Supplemental Fig. 6).

The average frequency of variant CpG sites occurring in a single DMR is 4.15%, 4.91% and 2.44% in inter-subspecies DMRs, intra-*Rnn* and intra-*Rnc* DMRs, respectively (Supplemental Table 7). In total, when calculated using the number of variant CpG sites in each DMR, 70.1%, 85.3% and 91.5% of inter-subspecies, intra-*Rnn* and intra-*Rnc* DMRs respectively carry no more than one variant CpG site and 93.8%-99.1% of the 3 DMR datasets carry no more than three variant CpG sites (Supplemental Fig. 5B). According to the definition of DMRs and average CpG number of a single DMR, such low variation in CpG sites within DMRs indicates that the DNA methylation level variation is unlikely directly caused by the CpG variation, but driven by other forces.

### Discussion and conclusion

The results from phylogenetic inference, Tajima’s D calculations and the ratio of SS-SNVs all indicate strong selective signals within the inter-subspecies DMRs but not intra-subspecies DMRs or non-DMRs. Both inter-subspecies DMRs and intra-subspecies DMRs have higher nucleotide diversity than non-DMRs, however, only inter-subspecies DMRs show accelerated fixation of SNVs. Thus, accelerated DNA evolution in inter-subspecies DMRs is comprised of two independent processes: increased nucleotide diversity and accelerated fixation of these DNA variants.

Deamination of methylated cytosines has been suggested as the main mechanism of nucleotide diversity variation across the genome (Fryxell and Moon 2005; Mugal and Ellegren 2011; Xia et al. 2012; Zhao and Jiang 2007). Our results demonstrate that increased nucleotide diversity in DMRs is driven by variation of methylation level rather than deamination of methylated cytosines, indicating changed epigenetic status could be the root cause of increased nucleotide diversity in DMRs. A recent review suggested that methylation changes occur downstream of gene regulation during cellular differentiation (Baubec and Schubeler 2014). Similarly, gene expression changes are necessary for methylation changes to occur in response to environmental shifts. This lets us speculate that it is adaptation to the changing environment leading to changes of epigenetic status, which in turn induce the increased nucleotide diversity. A recent study reported that replication defects, which result from chromatin changes caused by a DNMT3B mutation, can cause differences in individuals with mutations (Lana et al. 2012), indicating a correlation between methylation status and mutation rates. CpG content dependent correlation between non-CpG and CpG mutations (with a threshold of ~0.53% CpG content) (Walser et al. 2008; Walser and Furano 2010) also supports a role for methylation in DNA mutation. Epigenetic status may have an impact on the fidelity of DNA replication, causing an increased replication error rate (Loeb and Monnat 2008; McCulloch and Kunkel 2008; Walser et al. 2008; Walser and Furano 2010). This implies that replication could be affected by chromatin changes. Germ line DNA mutations occur mainly during replication in meiosis. As reviewed in previous studies, alteration of the sperm epigenome, including DNA methylation, histone modification and sRNA have now been shown to be a mechanism of transgenerational inheritance (Boffelli and Martin 2012; Heard and Martienssen 2014; Lim and Brunet 2013), and, these changes do affect chromatin status. Additionally, reprogramming finishes during meiosis (Seisenberger et al. 2013), and methylation status is faithfully maintained during DNA replication in meiosis. Thus, these data support a model in which adaptation to the changing environment leads to changes in epigenetic status, which in turn induces higher nucleotide diversity (Fig. 6).

**Figure 6.**
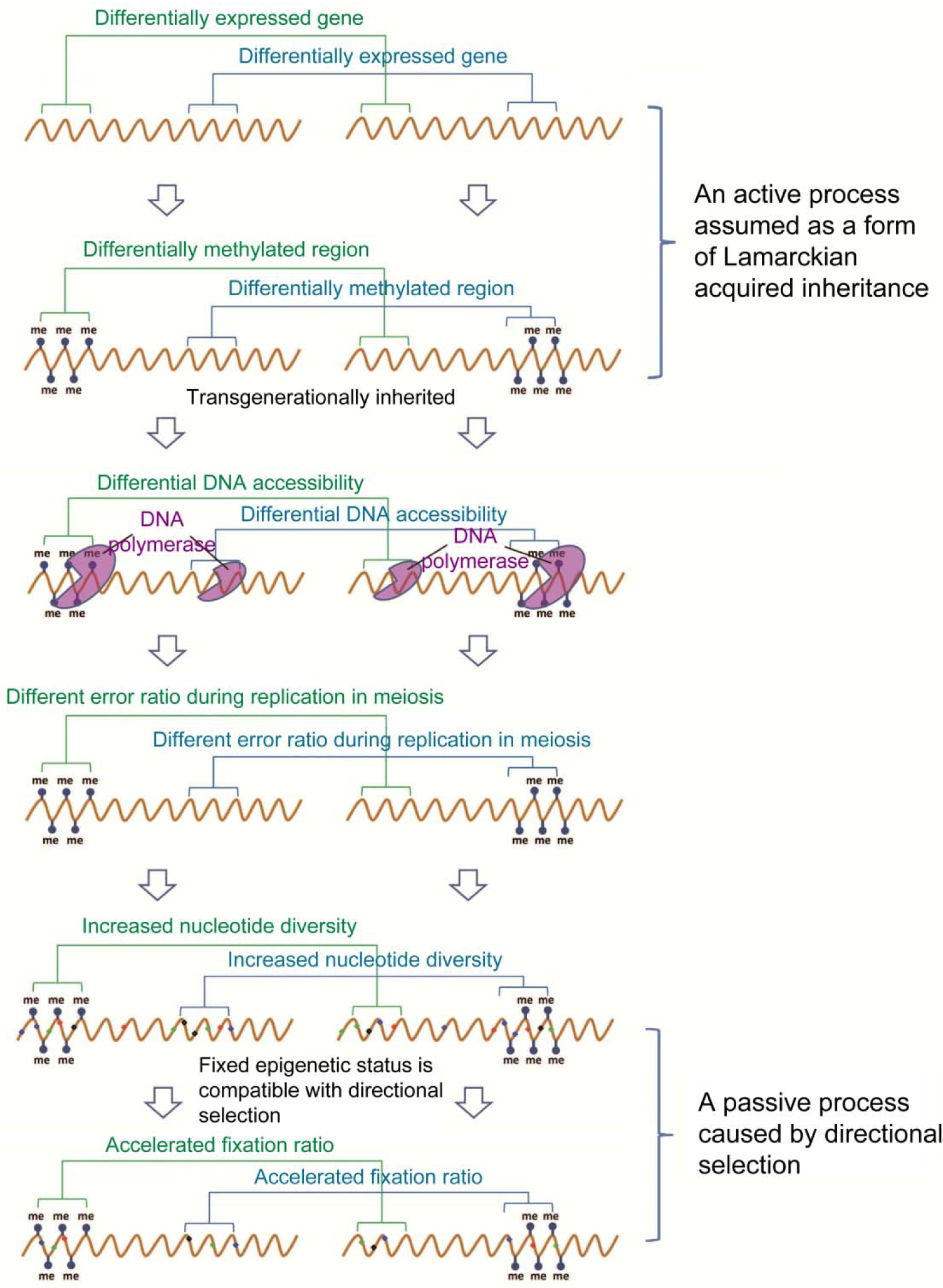
Hypothesized model of increased nucleotide diversity and accelerated fixation ratio in DMRs. Schematic of the model in which environmentally induced methylation differences lead to increased DNA substitutions and accelerated fixation. Orange lines represent DNA, and colored dots on the DNA represent substitutions.

An alternative hypothesis is that the DNA sequence in DMRs may be mutation hotspots, which may drive compatible variation of methylation status as suggested previously (Liu et al. 2014). However, three key pieces of evidence make this unlikely. First, we found significantly increased nucleotide diversity between subspecies but decreased nucleotide diversity within each subspecies in inter-subspecies DMRs. Second, in intra-*Rnn* DMRs, comparatively the majority of these regions are non-DMRs in *Rnc* (Supplemental Fig. 1), we only see increased nucleotide diversity between the two individuals of subspecies *Rnn* but not *Rnc*, and vice versa. Third, sampled non-DMRs with similar nucleotide composition but low CpG density indicate the strong relationship between increased nucleotide diversity and CpG density. However, CpG content is relatively more conserved in DMRs than in non-DMRs. Thus, we can conclude that it is variation of methylation level inducing changes of chromatin accessibility, which in turn leads to increased nucleotide diversity. Changing the underlying DNA sequence can be a slow process that isn’t dynamic enough to respond to rapid environmental change (Heard and Martienssen 2014), especially for complex traits. Logically, methylation changes caused by active regulation of gene expression in response to changing environment should occur prior to increased DNA variation, because DNA mutation is a random process that can’t support dynamic gene expression regulation.

Directional selection increases the fixation probability of DNA variants. However, accelerated fixation of DNA variants occurs only in inter-subspecies DMRs. Since both intra-subspecies DMRs (caused by random and temporary environmental changes) and inter-subspecies DMRs (caused by stable and long-standing environmental changes) show increased nucleotide diversity, this is unlikely to be a consequence of natural selection. However, increased nucleotide diversity may provide a substrate for natural selection to act upon.

The subspecies-specific methylation status in inter-subspecies DMRs implies that methylation statuses can be maintained by stable environment differences. Additionally, because we saw accelerated fixation in inter-subspecies DMRs but not intra-subspecies DMRs with similar nucleotide diversity, we speculated that stably diverged environments act as a kind of directional selection, leading to fixation of specific gene expression patterns and associated epigenetic status, which eventually fix advantageous DNA variants (Fig. 6).

Epigenetic inheritance can be compatible with Darwinian evolution if epigenetic statuses that specify traits can be transgenerationally inherited (Boffelli and Martin 2012). Our results demonstrate that not only environment-associated methylation variation can be maintained and transgenerationally inherited, but also lead to increasing of nucleotide diversity. We show that fixed methylation status is compatible with Darwinian evolution, and can lead to accelerated fixation of DNA variants that confer an advantage in the new environment. These results establish a bridge between Lamarckian acquired inheritance and Darwinian selection.

## Methods

### Experimental Paradigm

The two selected subspecies, *R. n. caraco* and *R. n. norvegicus*, live in extremely different environments from north and south China, respectively. Harbin City has a severe winter (126°32E, 45°48′N) compared to Zhanjiang City (110°21E, 21°16′N), which is the main reason for reproduction inhibition in the subspecies *R. n. caraco*. Two individuals were selected for each subspecies, and both the genome and the sperm methylome were sequenced for each sample. We proofread the methylomes using the genome sequence of each individual to ensure the statistical reliability of relationship between methylation and DNA variation.

We examined DMRs with methylation level changes greater than two fold and methylation level differences (MLD) greater than or equal to 0.2 in the methylomes between the two subspecies (inter-subspecies DMRs), or within *R. n. caraco* (intra-*Rnc* DMRs) and *R. n. norvegicus* (intra-*Rnn* DMRs), respectively. Inter-subspecies DMRs are defined as DNA regions with significantly higher inter-subspecies MLD than intra-subspecies MLD (see **Differential Methylation Analysis**). The genomic regions excluding the inter- and intra-subspecies DMRs are defined as non-DMRs. Genome sequences were classified as DMRs and non-DMRs for further analysis.

Because nucleotide diversity (*π*) and other indices of variation are sensitive to the length of DNA sequences, two methods were used for sampling of non-DMRs as follows: First, non-DMRs with equal length and GC content to each unique inter-subspecies DMR set are randomly sampled, and then the sampled 1859 non-DMRs are combined into one sequence, with length equal to the total length of inter-subspecies DMRs (2.6Mb). This sampling process is repeated 1,000 times, and decreases the influence of sequence length on π and other statistics. Second, because the mean length of DMRs is less than 1.5kb, non-DMRs with equal length and GC content to each unique DMR are randomly sampled 10,000 times from the genome, and the mean π of these sampled non-DMRs is used as the value of the non-DMR corresponding to that unique DMR to reduce the randomness and bias of sampled non-DMRs. This method enables comparison of the landscape of variation in a single DMR within the three DMR datasets.

### Sperm Collection and DNA extraction

All animal experiments were conducted with the permission of the Institutional Animal Use and Care Committee of the Institute of Plant Protection, Chinese Academy of Agricultural Sciences.

Mature sperm were isolated from cauda epididymides as described by Kempinas with some modifications (Kempinas and Lamano-Carvalho 1988). The cauda epididymidis was cut longitudinally in a 35mm diameter Petri dish with 3ml M2 medium (M7167, Sigma). The sperm were released by repeatedly and gently disrupting using a pipette tip after incubation at room temperature for 20min, and were then collected and washed twice in PBS. To eliminate somatic cell contamination, the sperm were treated with somatic cell lysis buffer (0.1% SDS, 0.5% Triton X in DEPC H2O) for 20 min on ice (Peng et al. 2012). Purified sperm were then incubated in SNET buffer (20 mM Tris-HCl, 5 mM EDTA, 400 mM NaCl, 1% (wt/vol) SDS; pH 8.0) containing 400µg/ml Proteinase K and 40mM DTT in a 55°C shaker (~ 150) overnight. DNA was harvested using a standard phenol-chloroform extraction protocol. For extraction of DNA from testis, the SNET buffer with only Proteinase K was used, followed by the standard extraction protocol. DNA was eluted in TE for sequencing.

### DNA Isolation, BS-Seq Library Construction and Sequencing

Five µ genomic DNA was first fragmented by sonication with a Covaris S2 system (Covaris, MA) to a mean size of approximately 250 bp, followed by end repair, 3’-end addition of dA, and adapter ligation. Methylated adapters were used according to the manufacturer’s instructions (Illumina). The bisulfite conversion of sample DNA was carried out using a modified NH4HSO3-based protocol 1 and amplified with 9 cycles of PCR. Paired-end sequencing was carried out using an Illumina HiSeq 2000.

### Genome re-sequencing and BS-Seq Analysis

The *Rattus norvegicus* reference genome was downloaded from Ensemble (release-70). To avoid the failure of reads mapping caused by additional mismatches resulting from C to T transitions after bisulfite treatment, all Cs in the reference genome were converted to Ts (T-genome) and all Gs were converted to As (A-genome) separately, creating two reference genomes. Moreover, the sequenced reads were prepared for alignment by replacing observed Cs on the forward read with Ts and observed Gs on the reverse reads with As. We used SOAP2 (Version 2.21) (Li et al. 2008; Li et al. 2009) to map the transformed reads to both the T- and A- genomes, allowing up to 6 mismatches in 90 bp paired-end reads. Reads aligning to more than one position on the genome were discarded. Multiple reads mapping to the same position were regarded as PCR duplicates, and only one of them was kept. For mC detection, we retrieved the original sequence of the transformed reads and compared it with the untrasformed reference genomes. Cytosines in BS-seq reads that matched to Cs on the reference were counted as potential mCs. Cytosines with a quality score < 20 were not considered.

The bilsulfite conversion rate of each library was calculated as the total number of sequenced Cs divided by the total sequencing depth for sites corresponding to Cs in non-CpG sites. The conversion rates of all the libraries was higher than 99%. To distinguish true positives from false positives, we used a model based on the binomial distribution B(n,p), with p equal to the false positive rate and n equal to the coverage depth of each potential mC. For example, given a potential mC position with k sequenced cytosines and total depth of n, we calculated the probability that all the k cytosines sequenced out of n trials were false positives, and then compared the probability of B(k,n,p) to 0.01 after adjusting p-values by the FDR method (Benjamini et al. 2001). Only the mCs with adjusted *p*-values < 0.01 were considered true positives.

### Methylation Level Calculation

The methylation level of an individual cytosine was determined by the number of reads containing a C at the site of interest divided by the total number of reads containing the site. Methylation level of a specific region was determined by the sum of methylation levels of individual cytosines in the region divided by the total number of covered cytosines in this region.

### Differential Methylation Analysis

Two-way analysis of variance (two-way ANOVA) was conducted to identify differentially methylated regions (DMRs) between two groups of samples (i.e. G1+G2 vs H1+H2) using 200, 500, 800, 1,000, 2,000, 3,000, 4,000, 5,000, 10,000 bp sliding windows with step lengths of 50% of the window size. As methylation at CpG sites is symmetric, we combined the data from the plus and minus strands for each CpG site during DMR detection. To ensure adequate power in the statistical test, only windows with at least 6 informative CpGs (≥ 5X coverage) in all four sequenced samples were considered. The two independent variables for ANOVA were group and cytosine position. For each window, we first calculated the variance between groups (variance caused by inter-group differences) and the variance between two individuals within the same group (variance caused by inter-individual differences), then used an F-test to calculate the *p*-value of each window by comparing the inter-group variance and inter-individual variance. *P*-values were then adjusted for multiple testing by the FDR method (Benjamini et al. 2001). Only windows with adjusted *p*-value < 0.05 and > 2-fold methylation level change were considered as candidate DMRs. In addition, we removed all DMRs in which the differences in methylation levels were < 0.2. Finally, contiguous DMRs and DMRs identified with different window sizes were merged.

### Normalization Procedures for Rates of Each Substitution Type

The substitution rates of any SNV site was normalized as follows:

C/T site as example:

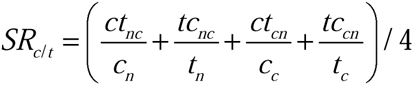

The *SR_c/t_* is the substitution rate of C/T sites. The *ct_nc_* is the total number of C to T substitutions when comparing the genome of *Rnn* to genome of *Rnc*. The *c_n_* is the total number of cytosines in the two *Rnn* genomes. The *tc_nc_* is the total number of T to C substitutions when comparing the genome of *Rnn* to genome of *Rnc*. The *t_n_* is the total number of thymines in the two *Rnn* genomes. The *ct_nc_* is the total number of C to T substitution reading from genome of *Rnc* to genome of *Rnn*. The *c_c_* is the total number of cytosines in the two *Rnc* genomes. The *ct_nc_* is the total number of T to C substitutions when comparing the genome of *Rnc* to genome of *Rnn*. The *t_c_* is the total number of thymines in the two *Rnc* genomes. Others and so on.

Substitution rates in CpG were normalized using the combined length of CpGs, and substitution rates of non-CpG were normalized using the length of non-CpG nucleotides.

### Population test of RSS

RSS (ratio of SS-SNVs) is the percentage of subspecies specific SNVs (SS-SNVs, Supplemental Fig. 3) among all SNVs. SS-SNVs was defined as sites with allele frequency from 0.25 – 0.5 (calculated by 8 chromosomes) and detected only in one of two subspecies but both two individuals. To increase the accuracy of RSS calculation, we added 10 genome sequences (5 for each subspecies, kindly provided by the Kunming Institute of Zoology, CAS), with about 3x coverage for further analysis. In the early study, we identified a total of 12,669,132 SNVs in the 4 genomes, among which, 3,236,117 (25.54%) are covered in the 14 genome data. For the 20,634 SNVs in inter-subspecies DMRs, 4300 (20.84%) are covered in the 14 genome data (Table 1).

We tested the allele distribution of each SNV site covered in the 14 genomes using the Chi-squared test with Yates’ continuity correction, and sites with *p* < 0.05 are defined as SS-SNVs. We further compared the difference of RSS between inter-subspecies DMRs and non-DMRs using the Chi-squared test with Yates’ continuity correction.

### Computational Methods

All DNA phylogenetic trees were constructed using the Neighbor-Joining method in the PHYLIP software package. Substitution model: Jukes-Cantor, Rates: uniform rates. We calculated nucleotide diversity using method described by Vernot et al (Vernot et al. 2012). Tajima’s D was calculated using program in software Mega-CC 6.0 (Kumar et al. 2012). CpG density was calculated as the number of CpGs/DMR length. Statistical analyses were performed using R package. The means plus or minus one standard deviation are reported unless otherwise noted.

### Data access

Data analyzed herein have been deposited in SRA with accession XXXXXXXX.

## Acknowledgements

We thank Joshua M. Akey from Department of Genome Sciences, University of Washington, USA for advice on genomic analysis strategies and explanation of results. We thank Rachel Gittelman and Wenqing Fu from the Department of Genome Sciences, University of Washington, USA, and Guoliang Wang from Institute of Plant Protection, Chinese Academy of Agricultural Sciences, China, for advice on manuscript preparation. We thank Dongdong Wu and Lin Zeng from Kunming Institute of Zoology, Chinese Academy of Science, China for providing additional 10 genome sequence data used for population test. This experiment was supported by The Agricultural Science and Technology Innovation Program, National Key Technology R&D Program (2012BAD19B02), and National Basic Research Program of China (973 Program, 2007CB109104).

## Author Contributions

XH.L. designed all experiments. XH.L., F.L., N.L., DW.W., and Y.S. prepared the figures and wrote the manuscript. XH.L., JM.L., QY.L., YB.J., and Y.S. performed genomic and statistical analysis. F.L., DW.W., N.L., and QY.L. contributed to design of the experiment. ZY.F., and L.C. contributed to the design of the project. DW.W., N.L., DD.Y., JJ.S., and L.C. performed sampling, sperm collection and other preparation of genome sequencing and BS-seq.

## Disclosure declaration

The funders had no role in study design, data collection, and analysis, decision to publish, or preparation of the manuscript. The authors declare no competing financial interests.

---

**Supplemental Figure 1.**
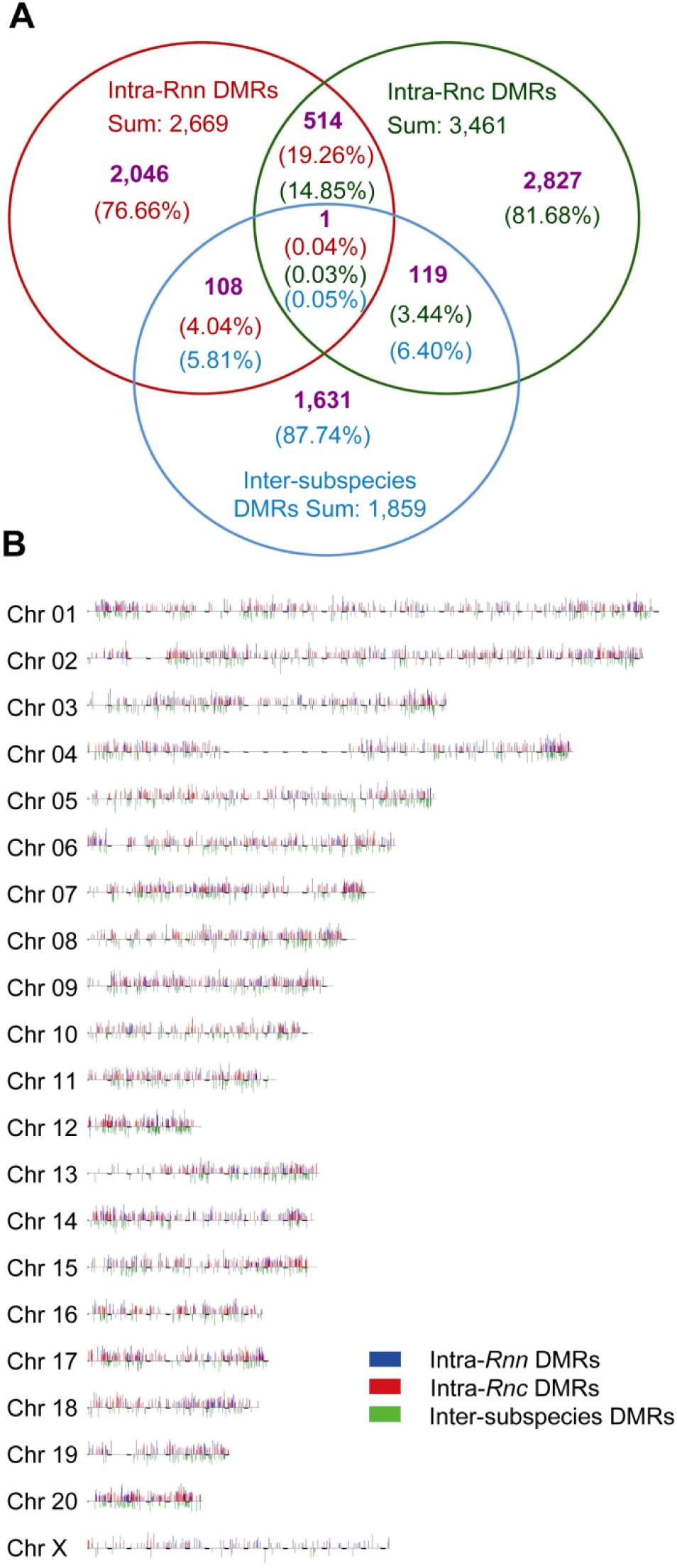
General features of the DMR landscape. (***A***) Identified DMRs and shared DMRs among inter-subspecies, intra-*Rnn* and intra-*Rnc* DMRs. See also Supplemental Table 2, 3, and 4. (***B***) Distribution of DMRs along the genome. Different DMR sets are illustrated with different colors. Positions of lines indicate the genomic locations of DMRs. The width of line is proportional to the length of the corresponding DMR, and the height of line is proportional to methylation level difference of the corresponding DMR.

**Supplemental Figure 2.**
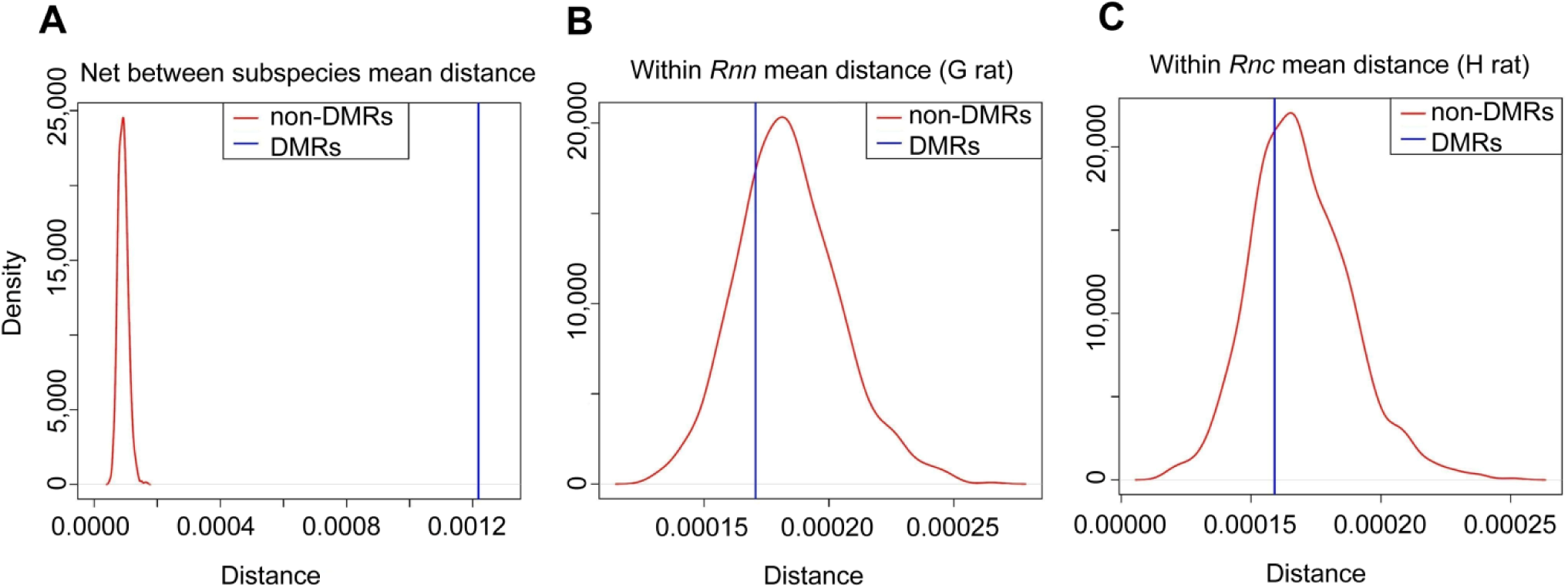
Comparisons of between and within group distances between intersubspecies DMRs and non-DMRs, Related to Fig. 1. **(*A*)** Comparison of between subspecies distance in inter-subspecies DMRs and non-DMRs. **(*B*)** Comparison of within *Rnn* distance (G1 vs G2) in inter-subspecies DMRs and non-DMRs. **(*C*)** Comparison of within *Rnc* distance (H1 vs H2) in inter-subspecies DMRs and non-DMRs.

**Supplemental Figure 3.**
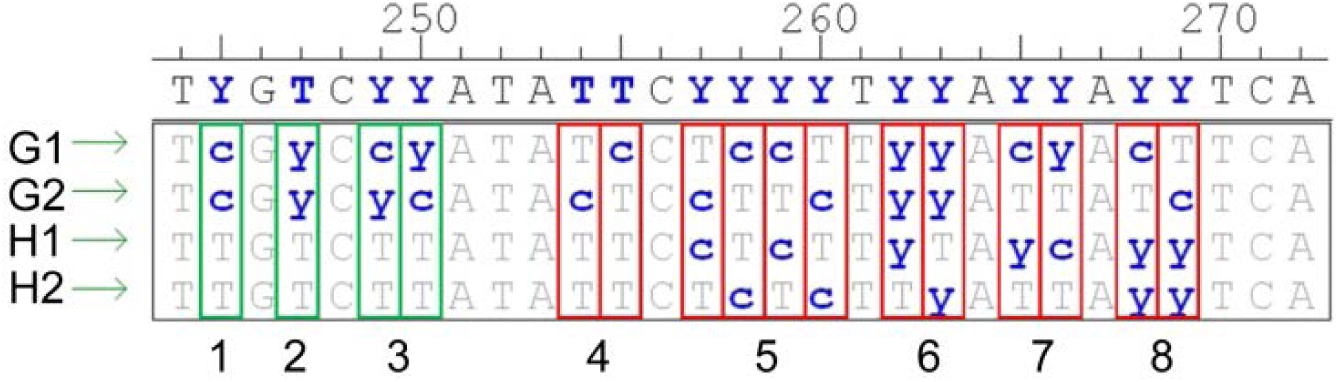
Illustration of subspecies specific SNVs, Related to Supplemental Experimental procedures. Because only two individuals of each subspecies were used to genome sequencing, we define **SS-SNVs** as sites with an allele frequency of 0.25 – 0.5 (calculated with 8 chromosomes) and detected in both individuals of one subspecies, but in neither individual of the second subspecies. Thus, **SS-SNVs** are sites as follows: Site1: The subspecies are fixed for two different alleles. Site 2: Both individuals are heterozygous in one subspecies, while in the other subspecies one allele is fixed. Site 3: One subspecies is fixed for an allele, while in the other subspecies one individual is homozygous for the other allele, and the other individual is heterozygous. In sites 4,5,6 and 7, there aren’t any variants that are private to one subspecies and present in both individuals, and they all are not **SS-SNVs**.

**Supplemental Figure 4.**
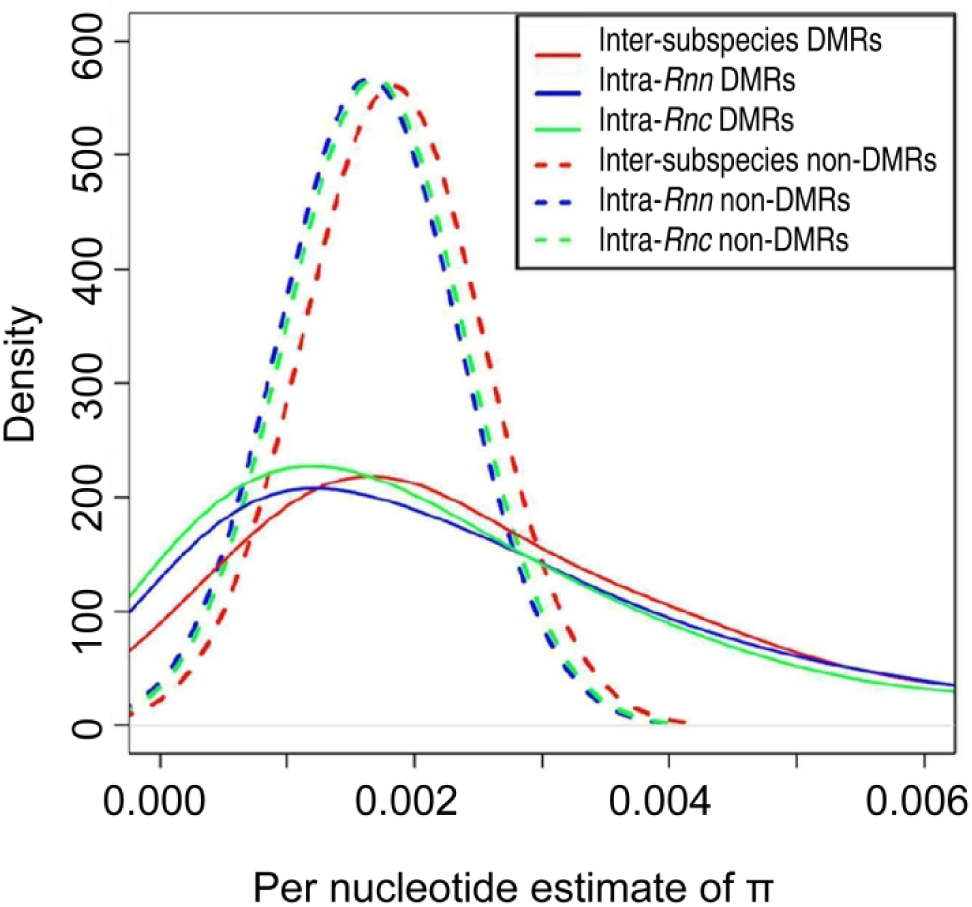
Distribution of π using the second sampling method. π of intersubspecies DMRs are calculated using 4 individuals, and π of intra-subspecies DMRs are calculated using 2 individuals of each subspecies respectively.

**Supplemental figure 5.**
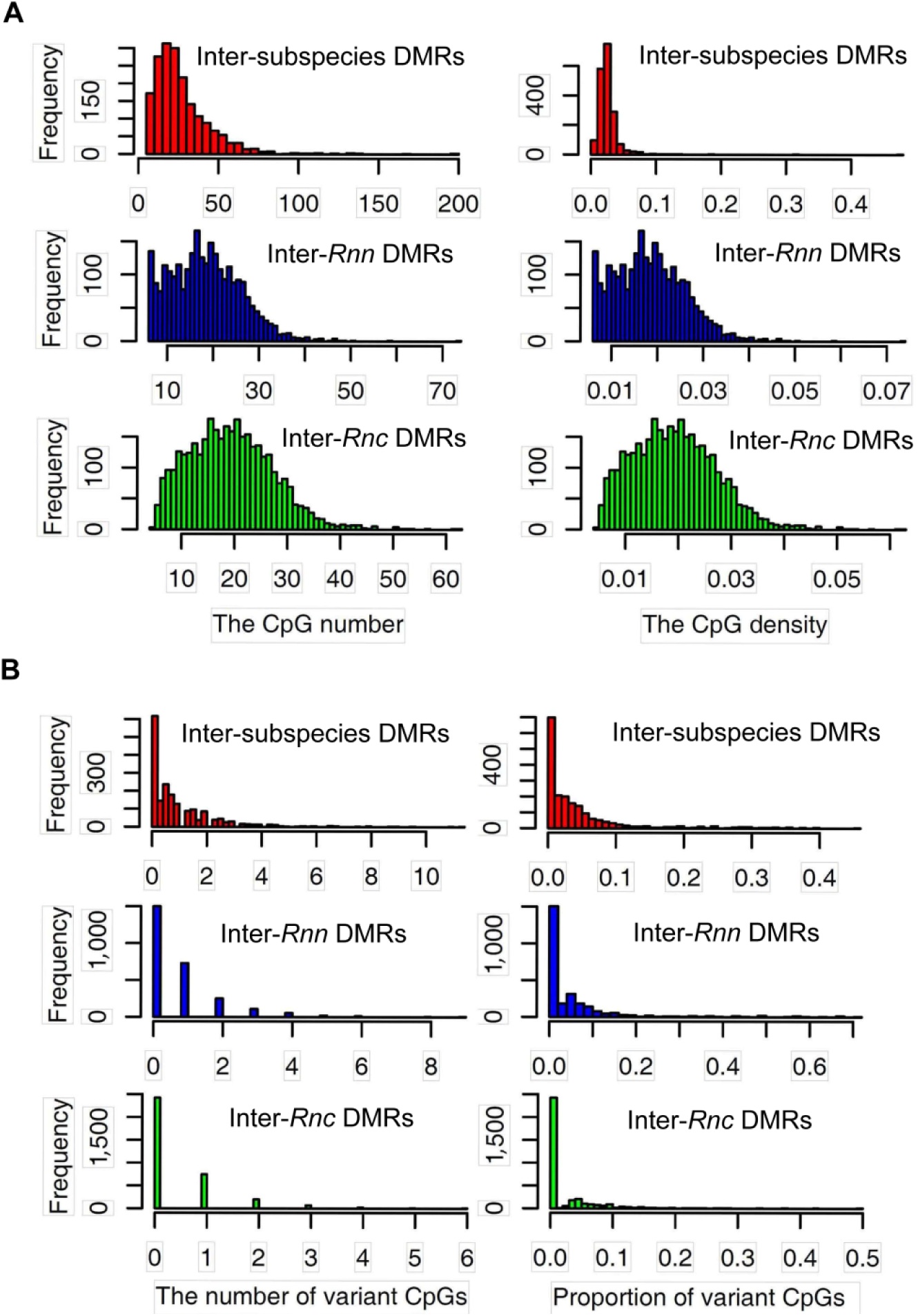
Characteristics of CpG content of DMRs. (A) Distribution of CpG numbers and densities within inter-subspecies DMRs, intra-*Rnn* DMRs and intra-*Rnc* DMRs. (B) Variation of CpGs in DMRs.

**Supplemental figure 6.**
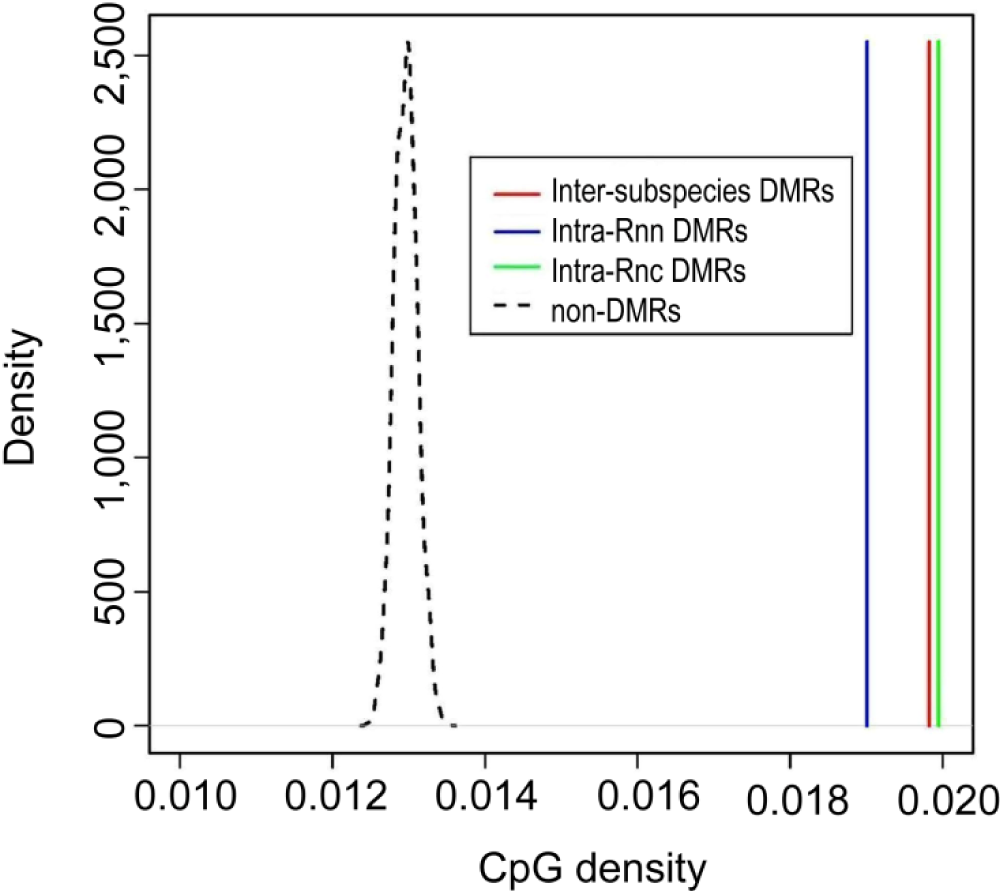
Comparison of CpG density. CpG density was calculated using the combined length of DMRs and corresponding non-DMRs using the second sampling method.

